# Ethnopharmacological disease classification and bioprospecting: the diversity of plant drugs used to treat cancer

**DOI:** 10.1101/2023.05.22.541754

**Authors:** Jamie Thompson, Julie Hawkins

## Abstract

Cancer is a highly-diverse disease and as the second-leading cause of death worldwide is a focus of drug discovery research. Natural products have been shown to be a useful source of novel molecules for the treatment of cancer. It is likely there are many plants with undiscovered molecules of therapeutic value, however identifying new leads from among the vast diversity of plants is very challenging. Traditional knowledge might inform bioprospecting by predicting lineages of plants rich in therapeutically useful molecules. Here, we characterise the phylogenetic diversity of plants used traditionally to manage cancer. We demonstrate the independent and repeated targeting of specific lineages of plants by different peoples in different parts of the world. That the same lineages are used to treat different cancers is suggestive of independent discovery of therapeutic value. However, the lineages we report here as rich in plants used traditionally to treat cancer coincide with those for other ethnobotanical applications, and contain few plants with proven anti-cancer activity. It is likely that the traditional knowledge recorded and explored here is shaped by selection of plants conferring milder effects for treating wider symptoms, such as tiredness or nausea, rather than for halting tumour growth. Accurate prediction of useful plant lineages for cancer management requires more nuanced information than is commonly provided in ethnobotanical records.

## Introduction

Cancer is a diverse set of diseases affecting different organs, unified by abnormal growth and division of cells into tumours (Melo et al., 2013; Taylor et al., 2013; Elia et al., 2015). It is the second leading cause of mortality worldwide, responsible for a sixth of deaths (WHO, 2018). While modern medicine relies on surgeries, radixo- and chemo-therapy for cancer management, a large fraction of the world’s population retains valuable traditional knowledge of plants used to manage cancer, which is well documented (Aumeeruddy and Mahomoodally, 2021). Many modern anticancer pharmaceuticals have been derived from plants (Spjut et al., 1976), including vinca alkaloids (Zhou and Rahmani, 1992), podophyllotoxin (Guerram et al., 2012) and taxanes (Shah et al., 2013). Correlative studies (Spjut and Perdue, 1976) and “reverse-ethnopharmacology” (Leonti et al., 2017) have strongly linked traditional knowledge to known cancer treatments, and those under clinical investigation. It is likely there are many plants with anti-cancer properties awaiting discovery, but these are difficult to identify. The contribution of traditional knowledge to modern medicine is vast, with an estimated ∼25% of pharmaceutical drugs derived from plants, and as many as 60% of antitumour drugs (Brower, 2008). Many more useful plants remain undiscovered, but only 15% have been evaluated pharmacologically (Verpoorte 1998; De Luca et al., 2012). The vast diversity of plants makes bioprospecting challenging, laborious and expensive (Firn 2003), but we can use traditional knowledge to improve efficiency (Cox and Balick,1994; Saslis-Lagoudakis et al., 2012; Halse-Gramkow et al., 2016). Characterising the phylogenetic diversity of plants used in local, folk medicine offers a method to expedite bioprospecting, by predicting lineages with elevated potential, so called “hot nodes” (Saslis-Lagoudakis et al., 2012; Halse-Gramkow et al., 2016). But this relies on the assumption that certain lineages used in traditional medicine are repeatedly targeted, and that their properties are useful for modern cancer medicine.

A large fraction of the world’s population in developing countries still rely on traditional medicine to meet healthcare needs (Twarog and Kapoor, 2004) and plants are a large component of this, with an estimated 10,000-53,000 species of plants used traditionally (FAO, 2003; McChesney et al., 2007). Understanding human-plant relations has been the focus of ethnobotanical research for centuries (Rahman et al., 2019; Balick and Cox, 2020), and is of growing practical importance. Traditional knowledge is being eroded through acculturation (Geck et al., 2016), language extinction (Cámara-Leret et al., 2021), and the rise of modern medicine. Similarly, plant diversity is being lost at an accelerated rate, largely due to human impacts (Antonelli et al., 2020; Nic Lughadha et al., 2020). It is vital not only to document traditional knowledge, but to understand patterns of medicinal plant selection (Souza et al., 2018; Teixidor-Toneu et al., 2018; Gaoue et al., 2021). Characterising the diversity of plants used in traditional medicine can not only refine understanding of important plant lineages, but also improves confidence in traditional health systems, and reveals forces shaping cultural knowledge (Saslis-Lagoudakis et al., 2014; Teixidor-Toneu et al., 2018, 2021b; Thompson et al., 2022).

Plants used in traditional medicine have sometimes been considered randomly selected, with little medicinal impact above placebo (Firezouli and Gori, 2007). But it has long been demonstrated that several lineages are selected preferentially (Molander et al., 2012; Saslis-Lagoudakis et al., 2012; Lei et al., 2020; Gras et al., 2021), probably due to discovery of lineage-specific phytochemistry. Phylogenetic comparative methods (PCMs) have been used to characterise patterns in the selection of medicinal plants in different contexts. Application of PCMs has revealed non-random lineage selection among distant cultures in specific genera (Saslis-Lagoudakis et al., 2011), entire ethnofloras (Saslis-Lagoudakis et al., 2011, 2012; Lei et al., 2020), and different aetiology systems (Lei et al., 2018). PCMs have revealed environmental and historical forces shaping knowledge (Saslis-Lagoudakis et al., 2014; Thompson et al., 2022), and non-random selection of plants targeting specific bodily systems including the nervous system and mind (Rønsted et al., 2012; Alrashedy and Molina, 2016; Halse-Gramkow et al., 2016). As well as providing evolutionary insights into ethnobotany, non-random selection strengthens support for efficacy of traditional knowledge and has been argued to indirectly evidence bioactivity (Saslis-Lagoudakis et al., 2011). Support is strengthened when lineages are discovered independently by distant populations, instead of via cross-cultural transmission of knowledge (Teixidor-Toneu et al., 2018). But many important diseases such as cancer have been neglected, despite comprehensive documentation of global traditional knowledge (Aumeeruddy and Mahomoodally, 2021).

PCMs could provide insight into whether specific plant lineages are preferentially selected for management of cancer, allowing prediction of lineages likely to harbour species with undiscovered utility, which has been demonstrated as a promising approach (Saslis-Lagoudakis et al., 2011, 2012; Ernst et al., 2016; Halse-Gramkow et al., 2016; Pellicer et al., 2018). But the utility of this approach in traditional cancer knowledge is unclear. Standardised medicinal use categories tend to be associated with bodily systems, and may not reflect underlying pharmacological action (Staub et al., 2015; Ernst et al., 2016). This can result in selection of related lineages for use across disease classifications (Lei at el., 2020), making predictions of useful lineages misleading, depending on the level at which they are performed. Whether plants used across cancer types are selected from related lineages is hard to predict. The few systematic investigations of medicinal plants in treating specific disease targets have focussed on single organ systems (Rønsted et al., 2012; Alrashedy and Molina, 2016; Halse-Gramkow et al., 2016) or ailments with similar pathologies (Molander et al., 2012). Plants used for different cancers may be related due to shared underlying pathology, unrelated due to organ-specific needs, or even selected for non-tumour effects, such as improving general health symptoms. Investigating phylogenetic clustering between plants used traditionally for cancer and plants with unrelated ethnobotanical applications can improve predictions of useful lineages. Plants used for cancer have been associated with plants used as poisons or antifertility treatments (Leonti et al., 2017), possibly because of the shared underlying property of cytotoxicity. However, the efficacy of traditional knowledge in identifying and directly treating tumours is questionable, given the sophisticated diagnostic technology required for modern medicine to do so. It is likely that traditional knowledge systems address more-obvious general-health symptoms associated with cancer, such as weakness, lowered immunity and nausea.

Here, we assess phylogenetic patterns of global traditional cancer knowledge in angiosperms using a comprehensive genus-level phylogeny, and a dataset of 597 genera used to manage cancer. We reveal non-random selection of plants used against cancer, and for most organ-specific cancers, and assess the possibility of predicting undiscovered useful lineages. Extensive cross-predictivity in plants used across cancer types reduces our ability to predict useful lineages for specific organs, but prediction is possible when considering cancer plants as a whole. However, further comparisons with plants undergoing clinical trials for cancer, and for unrelated ethnobotanical uses, suggests that many traditionally-used plants confer mild effects which are likely to target symptoms associated with cancer. Our results caution that detailed investigations are needed when informing cancer bioprospecting with traditional knowledge.

## Results

### Non-random selection of lineages used in traditional cancer management

Plants used in traditional cancer management overall (all types of cancer) are selected from related higher taxonomic lineages within angiosperms (NRI 9.09, p < 0.05), and the subset of ∼1,600 angiosperm genera with medicinal use reported by Mabberley (2017) (NRI 3.32, p < 0.05) (Figure 1). Similarly, plants used to manage specific cancers are significantly selected from related deep lineages (NRI > 1.96, p < 0.05), with the exception of plants used to manage colorectal, liver and stomach cancers, which show neither clustering nor overdispersion (NRI >-1.96 and <1.96, p > 0.05) (Table 1).

**Figure 1:**
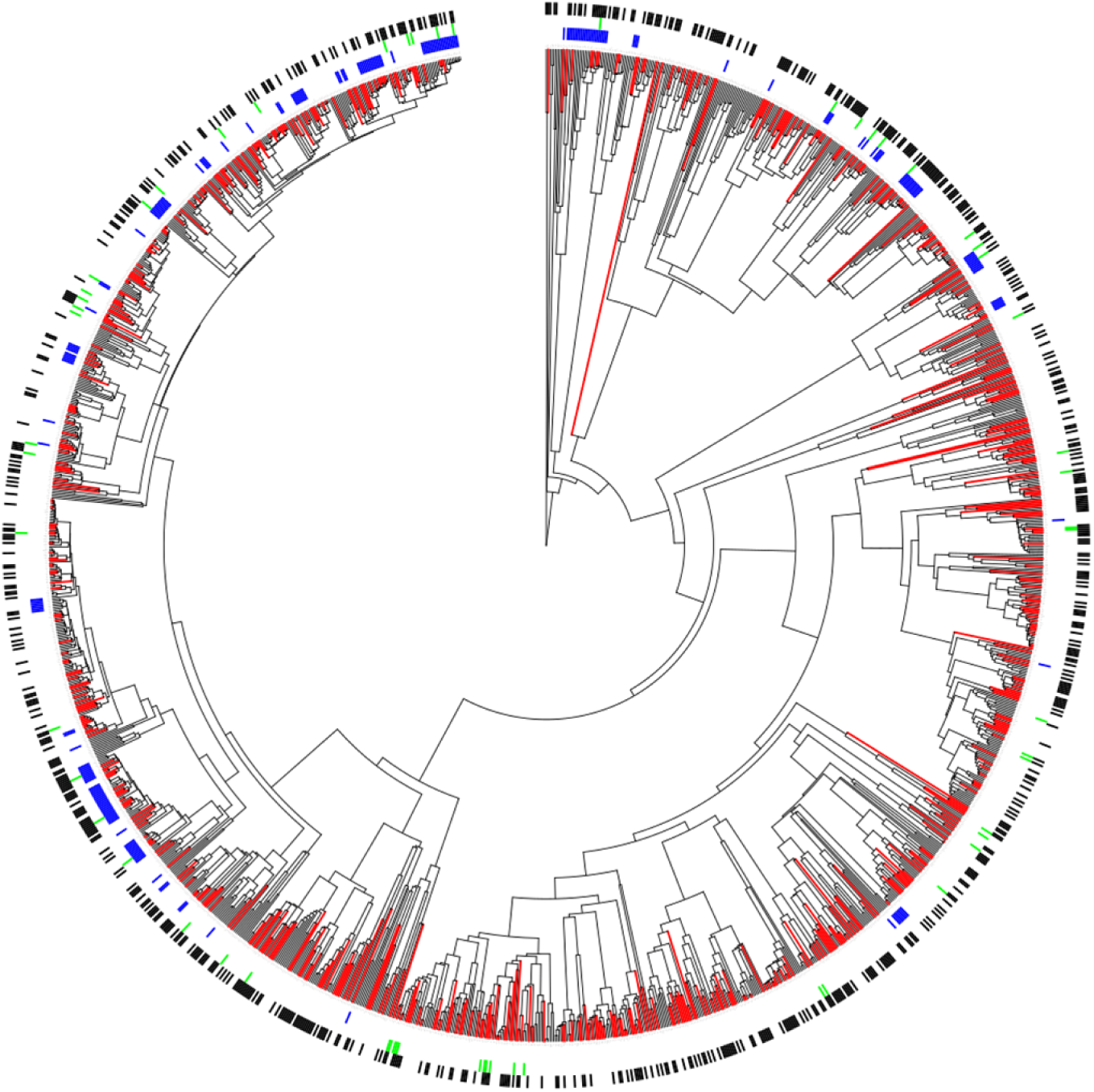
Phylogenetic distributions of genera used in traditional cancer management, in clinical trials for cancer, and edible plants plotted alongside lineages with predicted utility for bioprospecting. Genera in traditional cancer management (red branches) are phylogenetically clustered within medicinal plants and across angiosperms generally. Hot node lineages for ‘cancer genera’ (blue bars) are shown, as are genera including plants in clinical trials for therapeutic use as cancer drugs (green bars) and genera including plants with traditional use as food or food additives (black bars) are indicated. ∼19% of trials genera lie inside of hot nodes (13/67), and a similar proportion of food plant genera are within hot nodes (176/1,332; 13%), suggesting different properties shape traditional cancer knowledge. For ease of interpretation, we pruned genera without known medicinal use as described by Mabberley (2017) from the angiosperm tree in this figure, but analyses use all angiosperm genera unless specified otherwise.

**Table 1:** Phylogenetic clustering of genera used in traditional cancer management. Mean phylogenetic distance is calculated and compared with a null model of 9,999 random draws from the phylogeny pool, and significance is indicated with an asterisk. Clustering is assessed for genera with use across all cancer types within angiosperms generally, and within medicinal genera as described by Mabberley (2017). Clustering of genera used for specific cancer types is assessed within angiosperms.

| Target | Number of genera | Tree | NRI | Pattern |
| --- | --- | --- | --- | --- |
| All cancers | 597 | Angiosperm genera | 9.09* | Clustered |
| All cancers | 597 | Pruned to include only medicinal genera | 3.32* | Clustered |
| Breast | 165 | Angiosperm genera | 5.30* | Clustered |
| Cervical | 19 | Angiosperm genera | 2.5* | Clustered |
| Colorectal | 46 | Angiosperm genera | 1.68 | No pattern |
| Liver | 26 | Angiosperm genera | 0.30 | No pattern |
| Lung | 59 | Angiosperm genera | 4.33* | Clustered |
| Prostate | 56 | Angiosperm genera | 2.92* | Clustered |
| Skin | 137 | Angiosperm genera | 4.56* | Clustered |
| Stomach | 72 | Angiosperm | 1.88 | No pattern |
|  |  | genera |  |  |
| Throat | 44 | Angiosperm genera | 2.15* | Clustered |
| Uterus | 36 | Angiosperm genera | 2.09* | Clustered |

### Related lineages are targeted for use across most cancer types

Pairwise comparisons of clustering between genera used for cancers of different organs reveal high levels of cross-predictivity (Figure 2). Generally, certain lineages are repeatedly targeted for management of different cancers (NRI > 1.96, p < 0.05), with the exception of most comparisons with liver cancers, for which no pattern is found.

**Figure 2:**
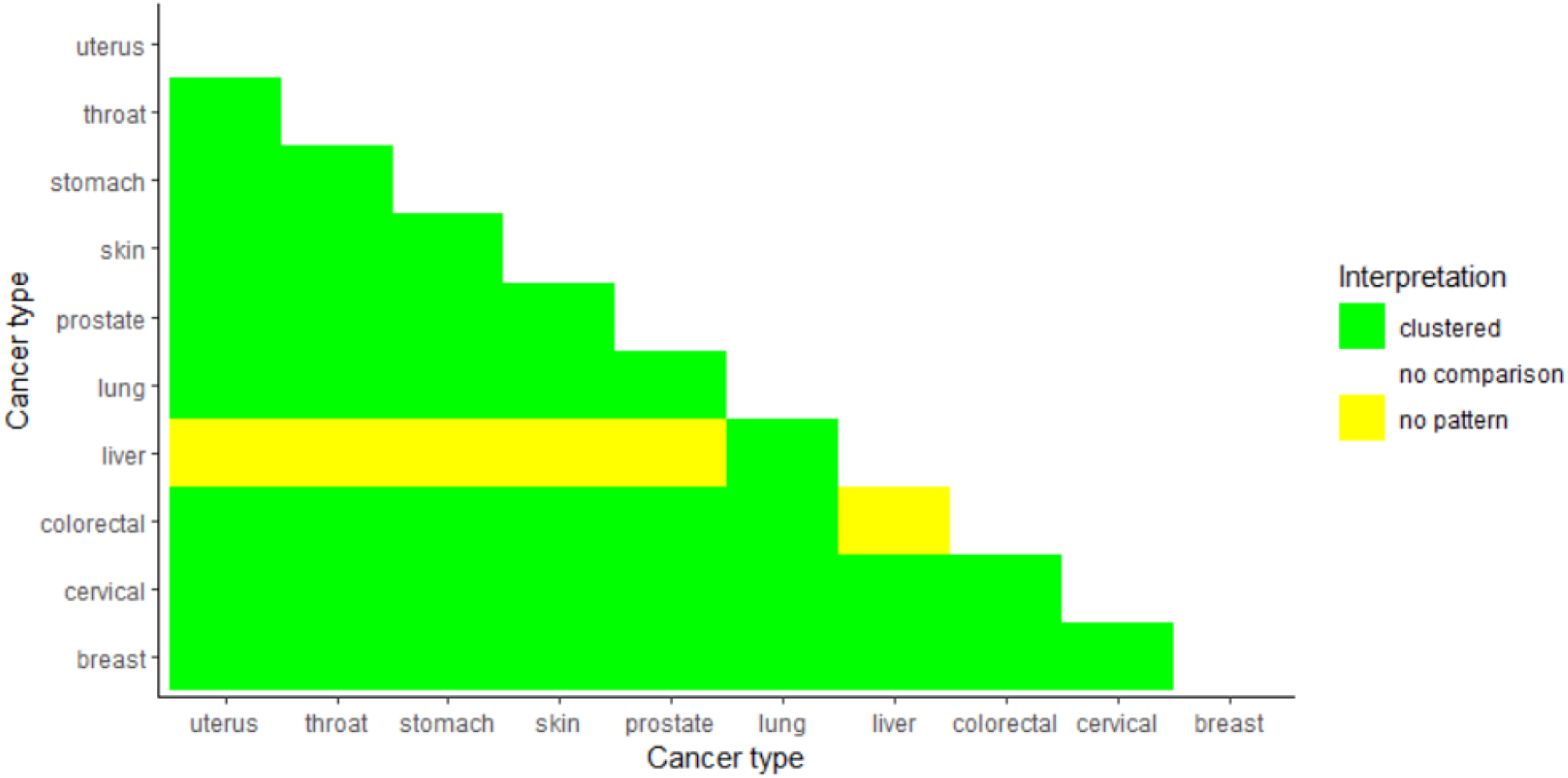
High cross-predictivity of lineages used for different cancer types reduces ability to predict organ-specific useful lineages. Genera used for specific cancers were selected from related lineages, except that only three of nine pairwise cross-comparisons that included genera used for liver cancer were significant. Significance is defined here as a net relatedness index (NRI) is greater than 1.96.

### Cross-predictivity between use in cancer and other ethnobotanical uses

Plants used traditionally for cancer management are selected from related deep lineages to those used for food, food additives, medicines, vertebrate poisons and invertebrate poisons (NRI > 1.96, p < 0.05), but not for antifertility uses (NRI >-1.96 and <1.96, p > 0.05) (Table 2).

**Table 2:** Plants used in traditional cancer management are drawn from deep lineages related to those of plants used for food and food additives, medicines across all therapeutic applications and poisons. They are not from lineages related to the lineages with antifertility uses.

| Ethnobotanical use | NRI | Interpretation |
| --- | --- | --- |
| Antifertility | 1.01 | No pattern |
| Food | 4.29* | Clustered |
| Food additives | 4.58* | Clustered |
| Invertebrate poisons | 5.37* | Clustered |
| Medicines | 9.58* | Clustered |
| Vertebrate poisons | 3.69* | Clustered |

### Independent discovery of useful lineages across continents, despite floristic variation

With the exception of Oceania, populations across continents select closely related deep lineages for cancer treatment (NRI > 1.96) (Figure 3a), but these are unrelated at shallower taxonomic depths (Figure 3b). At the shallower phylogenetic depth, several cross-continental comparisons are overdispersed (Africa and Asia, Africa and Europe, Asia and South America, Europe and North America) (NTI <-1.96), and no pattern is found for the remaining comparisons.

**Figure 3:**
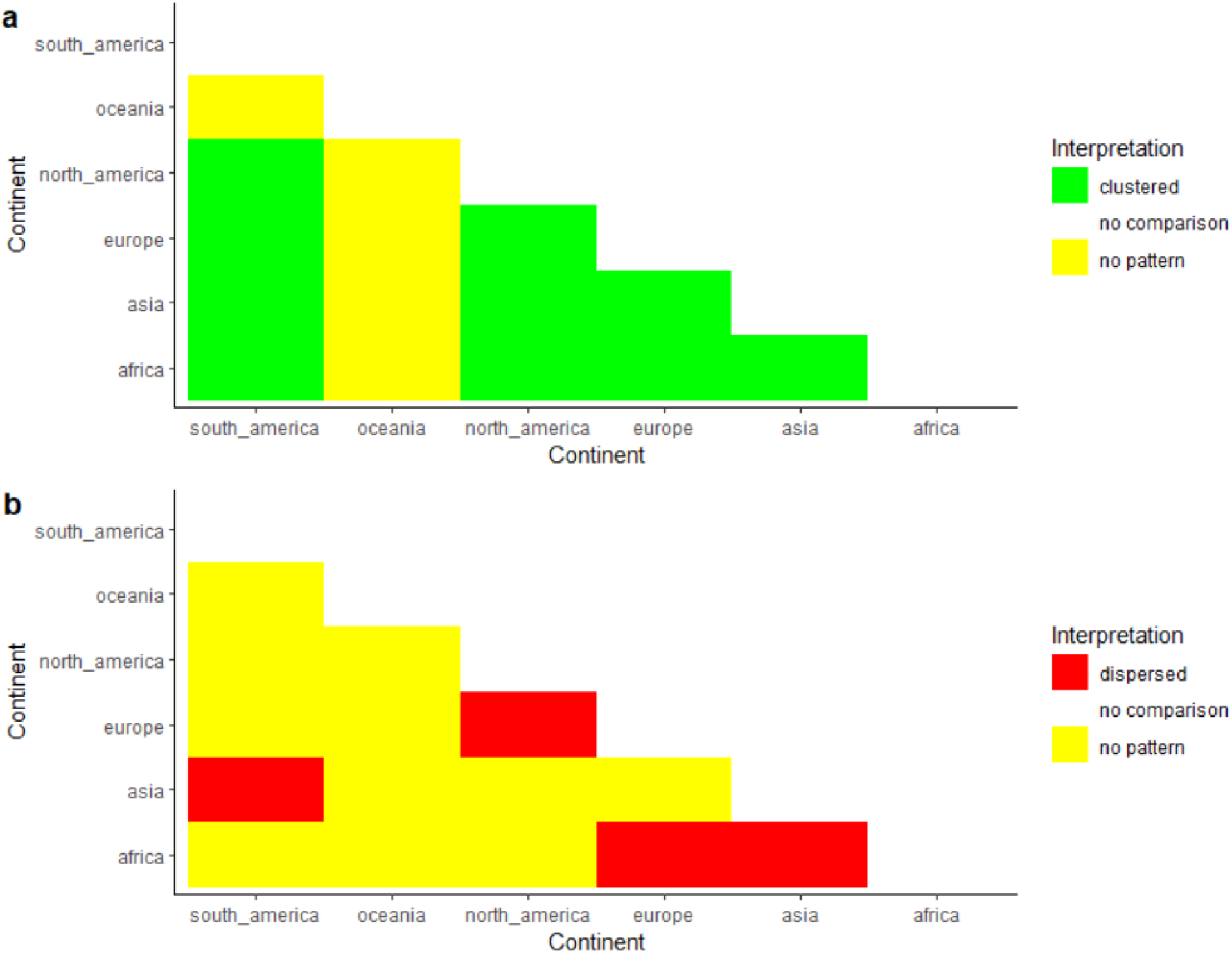
Distant populations select genera for traditional cancer management from related lineages (a), despite unrelatedness or even overdispersion at shallower taxonomic levels (b). Angiosperm floras differ across the globe, meaning that the same plants will not necessarily be available for selection.

### Predicting lineages with elevated bioprospecting potential

After confirming a high level of cross-predictivity in plants used for specific cancers, we predicted lineages with elevated utility against cancer generally. We identified 1,554 genera as having elevated likelihood of harbouring useful activity (∼13.36% of angiosperm genera sampled in the phylogeny), including 73 of the 597 cancer genera. These are distributed across 74 families, and the five families with highest representation are Apiaceae (263 genera), Lamiaceae (141), Rubiaceae (128), Araceceae (91), and Gesneriaceae (84). Two examples are plotted in Figure 4, demonstrating the predictive approach, and the full list of genera are given in Supplementary Materials. Of the 67 genera under investigation for cancer treatment in clinical trials (Zhu et al., 2011; Leonti et al., 2017), only 13 are found in the hot nodes (*Carum*, *Cocos*, *Curcuma*, *Cuscuta*, *Daucus*, *Drimia*, *Laurus*, *Mentha*, *Monarda*, *Panax*, *Papaver*, *Serenoa*, *Tabernaemontana*).

**Figure 4:**
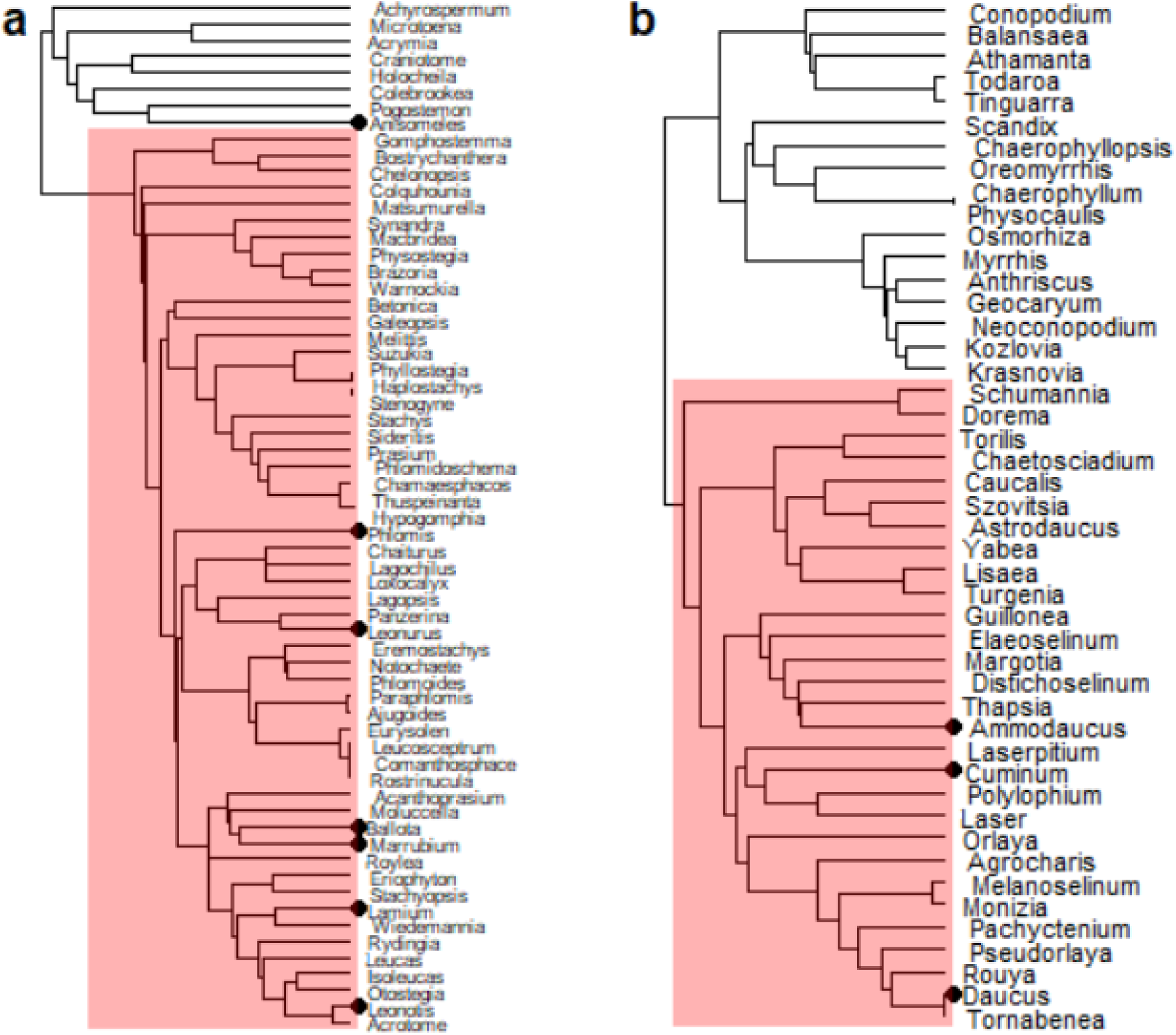
Two examples of taxonomically-unnamed lineages with elevated value for cancer bioprospecting, predicted by phylogenetic distributions of traditional knowledge. The predictive approach is demonstrated in two randomly selected cases. Genera with traditional use for managing cancer are denoted by a black circle at the tips, and the lineage with predicted elevated value is highlighted in red. Based on phylogenetic predictions alone, the red lineages should be prioritised in bioprospecting.

## Discussion

### Plants used for cancer treatments are non-randomly selected

Plants used traditionally for cancer management are phylogenetically clustered within angiosperms and within all medicinal plants (Figure 1, Table 1). The repeated targeting of certain lineages for therapeutic properties may indicate discovery of bioactive phytochemistry or useful pathways that are evolutionarily conserved (Saslis-Lagoudakis et al., 2011; 2012; Halse-Gramkow et al., 2016). When cross-cultural transmission of knowledge is low, similarities in medicinal knowledge can be interpreted as arising via independent discovery (Saslis-Lagoudakis et al., 2011, 2012; Hawkins and Teixidor, 2017; Teixidor-Toneu et al., 2018; Thompson et al., 2022). Independent discovery supports the view that plant medicines are efficacious (Bletter et al., 2007; Saslis-Lagoudakis et al., 2011, 2012). We found that plants used for cancer management are unrelated at lower taxonomic levels across continents, but are selected from related deep lineages, with the exception of Oceania (Figure 3). Relatedness at higher but not shallow evolutionary levels suggests selection of different plants from related, independently discovered lineages. Oceania holds knowledge not shared with other regions (Lloyd Jones and Sadgrove, 2015), likely explained by the evolutionary distinctiveness of the Oceanic flora (Carta et al., 2021).

### There are preferred plant lineages for treating most types of cancer

Considering plants used to treat each different cancer in turn, we showed that seven out of ten cancers were treated with plants that had a phylogenetic structure (Table 1). Lack of phylogenetic clustering of plants used against colorectal, liver and stomach cancers might be explained by cancer-specific properties. For instance, liver cancers are very complex pathologically, with multiple causes including toxin exposure, genetics, and hepatitis infections (Fan et al., 2013). Similarly, stomach cancers have complex causes including tobacco and infection by *Helicobacter pylori* (Balakrishnan et al., 2017). Variation in causes and symptoms may lead to selection of unrelated plants due to different lineage-specific properties. Several plants used against liver cancers are also used traditionally for hepatitis, with useful phytochemicals including oleanolic acid found in *Salvia*, and curcumin found in Zingiberaceae genera (Anand and Lal, 2016). Similarly, there is overlap between plants used traditionally against stomach cancers and for *H. pylori* infection, including *Alchornea*, *Allium*, *Calotropis* and *Terminalia* (Safavi et al., 2014). Presence of symptoms associated with comorbid infections may drive plant selection from unrelated lineages with different properties. A possible repercussion of the lack of clustering is that traditional knowledge is ineffective in identifying useful plant drugs for these organ-specific cancers.

### Related lineages of plants are used across cancer types

Whether selection is for cancers of specific organs, or the same plant drugs are used for different types of cancers, may be important for informing bioprospecting. A previous study showed the same plant lineages were used for the same specific therapeutic applications by people in different parts of the world, and this was interpreted as independent discovery of specific bioactivity (Saslis-Lagoudakis et al., 2012). This could suggest phylogenetic approaches based on specific therapeutic uses would be meaningful. However, recent research has cautioned against over-interpreting predictive patterns identified for specific therapeutic applications, because of high levels of cross-predictivity between therapeutic applications (Lei et al., 2020). We found that with the exception of plants used to treat liver cancers, related lineages are used to treat all cancers. Shared use of lineages across cancer types may be because therapeutic uses depend on general cytotoxic phytochemistry, targeting cellular processes underlying all cancer types perhaps by inducing cell death via inhibition of mitosis, DNA and ribosomal synthesis (Habli et al., 2017). Some of the most important plant-derived compounds used in clinical cancer medicine target multiple cancer types, including paclitaxel from the Pacific yew tree (*Taxus brevifolia*) (Priyadarshini and Keerthi, 2012), and vinblastine and vincristine, derived from the Madagascar periwinkle (*Catharanthus roseus*) (Mishra and Verma, 2017).

### Plants used against cancer include plants selected for cytotoxic properties

Leonti et al. (2017) used reverse ethnopharmacology to identify the traditional uses of plant drugs where proven value to biomedicine had been demonstrated. They found a statistically significant association between biomedical drugs with anticancer application and traditional uses for cancer therapy, but also a significant association between biomedical applications in cancer therapy and a subset of traditional gynaecological applications including abortifacients (Leonti et al., 2017). Leonti’s findings supported the observations of Spjut and Perdue (1976), that plants with uses as poisons were more likely to show cytotoxic effects. Later, phylogenetic study of the frequency of use of plants in ethnomedicine found that plants outside of the lineages that were frequently used appeared to have greater potential as leads in cancer medicine, as evidenced by frequency of clinical trials for plants belonging to and outside of hot nodes. Plants that lay outside of hot nodes, but were nevertheless used in ethnomedicine, were plants of infrequent use and strong effect and this too was attributed to the relationship between toxicity and potential as leads for development of anti-cancer drugs (Souza et al., 2018).

The distinction between plants used frequently for mild effects, and plants used infrequently for strong effects led us to explore whether there was cross-predictivity between uses for different therapeutic categories. We found clustering between plants used traditionally for vertebrate and invertebrate poisons and those used for cancer, suggesting that at least some of our hot nodes harbour cytotoxic phytochemicals which may be useful for cancer therapeutics. Support for the efficacy of traditional knowledge in identifying lineages with medicinal potential is strengthened by patterns of clustering between these two categories of use. Indeed, Mabberley (2017, categorised by Molina-Venegas et al., 2021) reported 292 genera including poisonous species; of these genera, 95 were amongst the 597 plant genera including species used for cancer treatment. However, whilst Leonti et al’s study (2017) might suggest clustering between plants used for anti-fertility and cancer treatment, we find no such relationship here. This might be because the anti-fertility category we use, from (Molina-Venegas et al., 2021), includes plants that are abortifacients but also contraceptives and fertility control, and is therefore a broad category.

### Plants used against cancer also include plants selected for properties unrelated to tumour growth

That the plants used in cancer treatment are significantly clustered with food plants indicates that properties unrelated to cytotoxicity are commonly selected for cancer management (Figure 1, Table 1). This result is not unexpected, because 328 of the 597 genera with plants used for cancer treatments in our study are among the 1,332 genera with uses as foods or food-additives, as described by (Molina-Venegas et al., 2021). The boundary between foods and medicines can be blurred, and the perception of medicines as food and foods as medicines is well-documented (Etkin and Ross, 1982, 1991; Johns 1990; Moerman 1994; Etkin 2008; Teixidor-Toneu et al., 2021a). The abundance of useful phytomolecules in food, such as antioxidants and nutrients, means edible plants are commonly used as medicines (Cisneros-Zevallos 2021). As many as one-third of plants used medicinally may be cultivated primarily as foods (Alqethami et al., 2017). Treatment of symptoms associated with cancer are likely treated with plants of mild effect, including food plants. Sufferers of cancer are commonly fatigued, with weakened immune systems. Food plants may be used medicinally to relieve these and other side effects. Many edible-plant derived antioxidants and anti-inflammatory compounds have been studied in modern biomedicine. These include curcumin in turmeric, lycopene in tomatoes and resveratrol in grapes (Russo et al., 2010).

### Can we identify lineages with elevated bioprospecting potential?

Lineages with higher proportions of medicinally-used taxa than expected by chance have been referred to as hot nodes, and several studies report hot nodes for medicinal plants overall or for specific therapeutic applications (Saslis-Lagoudakis et al., 2011, 2012; Halse-Gramkow et al., 2016; Ernst et al., 2018; Milliken et al., 2021; Cantwell-Jones et al., 2022). We estimated hot nodes, despite finding cross prediction between for specific cancer types, reasoning that these might be lineages with broad-spectrum anticancer properties. We identified ∼13.36% of angiosperm genera in 74 families with likely-elevated utility against cancer according to the hot nodes approach (Supplementary Materials). While this appears to offer a framework to improve future bioprospecting efforts, it is notable that just 13 of 67 genera (19.40%) with phytochemicals investigated in clinical trials for use against cancer are present in the hot nodes. If as previously discussed, therapeutic uses of plants in the treatment of cancer conflate gentle plants to support well-being and aggressive plants that might have anti-tumour activity, we might find a higher proportion of the plants in clinical trials in hot nodes that are significantly richer in plants used to treat cancer but which are not also food plants. The hot nodes reported here include 176 of 1,332 food-plant genera (13.21%); we caution that these plants may be a mixture of those selected for general symptoms and those with truly anti-tumour properties.

Our data suggest that plants used in traditional cancer management are diverse pharmacologically, and may encompass plants of mild effect for strengthening the patient, and plants with cytotoxic effect that might reduce cancer growth. The data that we used came from a secondary source that recorded type of cancer, but not the specific therapeutic goal of the plant drug intervention. Whether plants were used, for example, to manage nausea, was not recorded. Ethnobotanical data as reported in most publications is insufficiently nuanced to distinguish lineages likely to harbour anti-tumour phytochemicals, or have milder effect. Previous research into the utility of disease classification in traditional medicine has suggested that bodily categorisation performs poorly, and information on underlying biological responses are necessary (Ernst et al., 2016). Our findings similarly caution against interpreting classifications of therapeutic uses as pharmacologically meaningful. Where the goal of a bioprospecting endeavour is to identify potential anti-tumour compounds for cancer treatment, we recommend an ethnobotanically-informed phylogenetic exploration of plant poisons rather than one based on therapeutic application to treat cancer.

## Materials and Methods

### Data collection and processing

Genus-level data on flowering plants used in traditional cancer management were sourced from a recent and comprehensive systematic review, detailing 948 angiosperm species distributed in 153 families (Aumeeruddy and Mahomoodally, 2021). We recorded for each genus which cancer type it is used to manage, retaining only the top ten most common cancer types for the cancer-specific analyses (breast, cervix, colon, liver, lung, prostate, skin, stomach, throat and uterus), because the remaining 17 had too few plants for comparisons. No associations with cultures are provided, but country-level location data are described. We classified countries broadly into continents, which does not account for fine-scale effects of geographical proximity, but provides a broad test of whether distant populations select plants from related lineages, despite compositional differences of local floras. We did not consider migrant communities, because it is unclear whether they retained knowledge from their homelands, or adapted to the new region.

Genera with various relevant ethnobotanical uses were sourced from the fourth edition of the comprehensive and authoritative Mabberley’s plant book (Mabberley, 2017), compiled at genus-level by Molina-Venegas et al. (2021). In this compilation, genera are sorted in 28 use categories ranging from fuels and timber, to food and medicine. We retained six categories which may be associated with plants selected for cancer management (food and food additives, medicines, invertebrate and vertebrate poisons, and antifertility). Previous links have been made between cancer knowledge, poisons and antifertility drugs by Leonti et al. (2017). We additionally collected a list of genera in clinical trials for cancer (Zhu et al., 2011), compiled by Leonti et al. (2017).

## Phylogenetic analyses

### Testing for non-random selection of plants used in traditional cancer management

We used a large phylogeny of land plants sampling ∼13,000 genera in all analyses (Hinchliff and Smith, 2014), pruning non-angiosperm genera. We performed analyses of phylogenetic structure using the R package phylocomr (Ooms et al., 2023), which implements community phylogenetic methods available in Phylocom (Webb et al, 2008). We tested whether plants used in traditional cancer management for any cancer type were phylogenetically clustered at deeper taxonomic levels, by calculating MPD with the command “ph_comstruct”. The observed MPD was compared with 9,999 samples drawn randomly from across the phylogeny (null model 2), and the number of comparisons for which the observed distance was smaller or larger than the null samples was calculated. From this, two-tailed p-values and net relatedness index (NRI) were calculated. A positive NRI indicates clustering while negative indicates overdispersion, and significance is reached at >1.96 and <-19.6, respectively (at an alpha threshold of p<0.05 in a two-tailed p test). We tested for non-random selection by populations across the world despite geographic distance, cultural evolution and floristic differences among continents. This involved comparing NRI estimates of pairwise comparisons of plants used on each continent (estimated with the command “ph_comdist”) with the complementary metric nearest taxon index (NTI). NTI is calculated in the same manner as NTI using the command “ph_comdistnt”, but the metric mean nearest taxon distance (MNTD) is used instead of MPD. NTI brings insight into whether plants are selected from related lineages on a shallower taxonomic level. We assessed whether genera used in management of the ten best-reported individual cancer types are clustered at deeper taxonomic levels, with NRI.

### Testing for relationships between plants used for different cancer types, and with other ethnobotanical applications

We ran pairwise comparisons to understand the relatedness of genera selected for different cancer types, by calculating NRI between samples used for the top ten most frequently reported cancer types, using the command “ph_comdist”. We ran pairwise NRI comparisons between plants used against cancer and plants used for the six previously-described unrelated ethnobotanical uses (antifertility, food and food additives, vertebrate and invertebrate poisons, medicines).

### Prediction and description of lineages with elevated bioprospecting potential

We predicted hot nodes for cancer with the command “nodesigl” in Phylocom (Webb et al., 2008). This analysis identified lineages which were significantly overrepresented by genera used in traditional cancer management. To ensure we had effectively reduced the search for useful plants, we considered hot nodes which contained up to 100 tips, following Halse-Gramkow et al. (2016).

### Data visualisation

We visualised the phylogenetic distributions of traditionally-used plants with the Interactive Tree of Life v5 (Letunic and Bork, 2021). To visualise the two examples of predicted useful lineages, we used the R package ggtree (Yu et al., 2016). Heatmaps showing relatedness among different plant uses were produced with the R package ggplot2 (Wickham, 2011).

